# The Interplay of Bottom-Up Arousal and Attentional Capture during Auditory Scene Analysis: Evidence from Ocular Dynamics

**DOI:** 10.1101/2025.04.25.650619

**Authors:** Mert Huviyetli, Maria Chait

## Abstract

The auditory system plays a crucial role as the brain’s early warning system. Previous work has shown that the brain automatically monitors unfolding auditory scenes and rapidly detects new events. Here, we focus on understanding how automatic change detection interfaces with the networks that regulate arousal and attention, measuring pupil diameter (PD) as an indicator of listener arousal and microsaccades (MS) as an index of attentional sampling. Naive participants (N=36; both sexes) were exposed to artificial ‘scenes’ comprised of multiple concurrent streams of pure tones while their ocular activity was monitored. The scenes were categorized as REG or RND, featuring isochronous (regular) or random temporal structures in the tone streams, respectively. Previous work showed that listeners are sensitive to predictable scene structure and use this information to facilitate change processing. Scene changes were introduced by either adding or removing a single tone stream. Results revealed distinct patterns in the recruitment of arousal and attention during auditory scene analysis. PD was greater in REG scenes compared to RND, indicating heightened arousal in unpredictable contexts. However, no differences in overall MS activity were observed between scene types, suggesting no differences in attentional engagement. Scene changes—though unattended— elicited both PD and MS suppression, consistent with automatic attentional capture and increased arousal. Notably, only MS responses were modulated by scene regularity. This suggests that changes within predictable environments more effectively recruit attentional resources. Together, these findings offer novel insights into how automatic auditory scene analysis interacts with neural systems governing arousal and attention.

**Significance Statement:** Even without active listening, our brains automatically respond to changes in complex sound environments—like noticing a new sound on a busy street. These responses involve shifts in arousal and attention, helping us decide how to react, often without conscious awareness. Understanding this process is key to studying how we perceive sound scenes and how it may be disrupted in individuals with attention or arousal difficulties. In this study, participants passively listened to artificial soundscapes while we tracked eye activity: pupil dilation (a sign of arousal) and microsaccades (tiny eye movements linked to attention). We found that sudden scene changes triggered both responses, but they were differently influenced by scene predictability—suggesting they reflect separate aspects of automatic auditory processing.

## Introduction

The ability to rapidly respond to new events in our environment is crucial for survival. It is hypothesised that the auditory system functions as the brain’s “early warning system”, continuously monitoring the unfolding acoustic environment to quickly direct attention to new events (Cervantes Constantino et al., 2012; Murphy et al., 2013; Winkler and Denham, 2024). Indeed, listeners demonstrate a high sensitivity to abrupt changes—such as the appearance or disappearance of a sound source—even within complex, crowded auditory scenes (Eramudugolla et al., 2005; Pavani and Turatto, 2008; Cervantes Constantino et al., 2012; Puschmann et al., 2013; Petsas et al., 2016; Aman et al., 2021; de Kerangal et al., 2021). Notably, Brain responses, recorded from naïve distracted listeners indicate that such changes are often detected even in the absence of directed attention (Sohoglu and Chait, 2016a; b) supporting the notion that auditory change detection is, at least in part, an automatic process. A key factor enhancing this ability is the brain’s sensitivity to predictable structure in the environment. Change detection performance improves significantly when sound streams follow a temporally regular pattern compared to when they fluctuate randomly that this ability is enhanced by sensitivity to predictable structure in the environment (Aman et al., 2021; de Kerangal et al., 2021). Change detection performance improves when scene streams follow a temporally regular pattern rather than random fluctuations (Aman et al., 2021; de Kerangal et al., 2021). This finding is consistent with a broader literature indicating that the brain is highly attuned to statistical regularities in sensory input and leverages these patterns to facilitate more efficient interaction with the environment (Winkler et al., 2009; Bendixen et al., 2010; Bendixen, 2014; Barascud et al., 2016).

The general understanding emerging from these investigations suggests that the auditory system continuously and automatically monitors for changes in unfolding scenes, relaying this information to attention and arousal networks to initiate an appropriate response (e.g., fight or flight). In the present study, we sought to examine how this automatic change detection process interfaces with neural systems governing arousal and attention. To this end, we recorded eye and pupil dynamics while participants listened to ‘artificial acoustic scenes’ (see Methods). We analysed three types of ocular activity: pupil diameter (PD) and pupil dilation event rate (PDR), as indicators of arousal (Joshi and Gold, 2020), and microsaccades (MS), which serve as markers of attentional sampling (Pastukhov and Braun, 2010 ; Schneider et al., 2020; Zhao et al., 2024). Pupil diameter is a potential proxy for activity in the Locus Coeruleus (LC), which supplies the brain with the neurotransmitter norepinephrine (NE) that regulates vigilance and arousal. The LC exhibits two distinct modes of activation: tonic (sustained) activity, which is associated with general alertness and engagement, and phasic (transient) activity, which reflects rapid arousal responses to sudden or novel stimuli (Aston-Jones and Cohen, 2005; Joshi et al., 2016; Joshi and Gold, 2020; Wang and Munoz, 2021). Baseline pupil size is thought to reflect tonic LC activity, while brief dilations signal phasic responses.

Microsaccades (MS) are rapid, involuntary fixational eye movements hypothesized to reflect automatic exploration of the environment (Martinez-Conde et al., 2006; Otero-Millan et al., 2008). The presentation of novel stimuli triggers the temporary inhibition of MS (MSI) (Engbert and Kliegl, 2003; Rolfs et al., 2008; Wang and Munoz, 2021). Though MSI has long been thought to arise from a primitive sensory circuit, recent evidence suggests that it is modulated by stimulus salience and attention (Zhao et al., 2019b) and its dynamics can predict visual ‘attentional blink’ effects (Roberts et al., 2019) and awareness (White and Rolfs, 2016), highlighting its role in a sophisticated, rapid attentional re-orienting system that allows the brain to pause ongoing processing to evaluate sudden changes and select an appropriate behavioural response.

Here we investigate how the emergence of novel auditory events within a scene activates these pupil- and MS-linked systems. Better insight into how the auditory system interfaces with the brain systems that control arousal and attention is critical for elucidating the mechanisms that support effective responses in high-stakes time-critical situations.

## Materials and Methods

### Ethics

The research was approved by the Research Ethics Committee of University College London. Participants provided written informed consent and were paid for their participation.

### Participants

36 paid participants (26 Female; mean age:23.5, range: 18-34, SD=4.58) were recruited. All reported normal hearing with no history of otological or neurological disorders and normal or corrected-to-normal vision, with SPH prescriptions no higher than 3.5. One participant was excluded from the pupil diameter and microsaccade analysis due to exceedingly long reaction times on the decoy task (see below); three participants were excluded from pupil diameter and microsaccade analysis because of difficulty tracking the eye or excessive blinking or tiredness; three participants were excluded from microsaccade data analysis due to non-availability of binocular recording (see below information on detecting microsaccadic activity).

### Stimuli

Stimuli (Figure 1) were essentially identical to those used in Aman et al. (2021), except they were made longer (9 sec long instead of 4 sec long) to accommodate the expected slower pupil responses. ‘Scenes’ were populated by 6 streams of pure tones, representing 6 concurrent sound sources. Each stream had a unique carrier frequency (chosen from a pool of nine fixed values between 500-3225 Hz, spaced 2 cams on the ERB scale; ‘Equivalent Rectangular Bandwidth’; Moore and Glasberg (1983) and a unique temporal structure. These ‘scenes’ were thus perceived as composite ‘soundscapes,’ in which individual streams are perceptually separable. As such, they serve as effective models for busy natural acoustic environments (Cervantes Constantino et al., 2012). In ‘regular’ (REG) scenes, the streams had a regular temporal structure: For each stream, tone pip duration and inter-tone-interval duration were each chosen randomly from between 10 ms and 150 ms and then fixed for the duration of the scene. This yielded ‘sources’ with a variety of rates (from 3-50Hz) spanning the range which characterizes speech (Rosen, 1992) and many natural sounds. In ‘random’ (RND) scenes, the streams had a random temporal structure: For each stream, tone pip duration was chosen randomly from the values above and fixed for the duration of the scene, but the inter-tone-interval varied (also chosen from the range above) to yield an irregular temporal pattern. All streams (in REG or RND scenes), were phase-randomized (such that they started with the tone or the silent ITI). Each tone was ramped on and off with a 3-ms raised cosine ramp.

**Figure 1:**
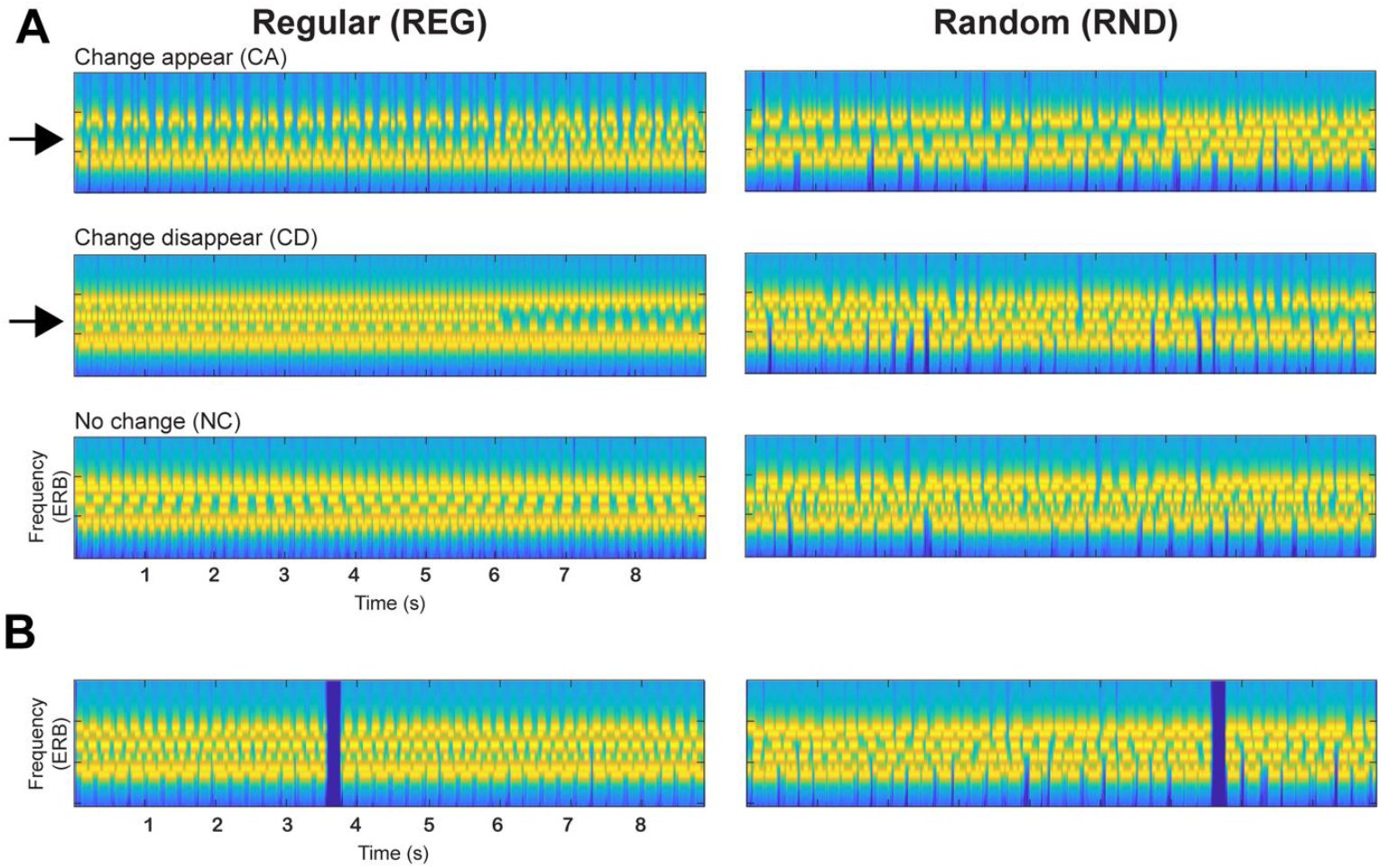
Change detection paradigm. **A:** Example of the three variations (‘change appear’, ‘change disappear’, and ‘no change’) of a scene with 6 streams. Regular (REG) scenes are on the left, and random (RND) scenes are on the right The changing component is indicated with an arrow. **B:** Examples of gap containing scenes.

Scenes in which each source is active throughout are referred to as ‘no change’ (NC) scenes. In ‘change appear’ (CA) scenes, a single stream is added partway through the scene.

In ‘change disappear’ (CD) scenes, a single stream is deleted partway (Figure 1). The change (appearing or disappearing) component was chosen randomly for each scene. The timing of the change in CA scenes, defined as the time at which the first tone pip of the appearing stream was presented, was 6 seconds after scene onset. For CD changes in REG scenes, the time of change was set to the offset of the last tone augmented by the inter-tone interval, i.e., at the expected onset of the next tone, which is the earliest time at which the disappearance is theoretically detectable. Therefore, change time varied somewhat from scene to scene but was always around 6 seconds post-onset. For CD changes in RND scenes, change time is ill-defined (because the temporal pattern is random). Hence, following the approach adopted in Aman et al. (2021) and de Kerangal et al. (2021), the change time in those scenes was set to the offset of the last tone augmented by the mean inter-tone interval (80ms).

The stimulus set also included ‘decoy’ (task-relevant) scenes (20% of the stimuli) that contained a 200 ms silent gap that the participants were instructed to detect and respond to as soon as possible. The gap was inserted randomly anywhere between 1s post onset to 1s pre offset. The task served the purpose of keeping the participants alert and broadly engaged with the auditory stimuli but was calibrated to be easy so that it minimally draws on attention/computational resources.

Overall, the main experiment consisted of eight ∼7 min-long blocks, each containing 30 trials: 4 trials of each of the main conditions REG-CA, REG-CD, REG-NC, RND-CA, RND-CD, RND-NC and 1 trial of each condition containing a gap. The stimuli were presented in random order. In total, 32 trials of each of the main conditions were available for analysis. The decoy (gap-containing) trials (48 overall) or any other trials that contained a button press (‘false alarm’) were not included in the analysis of the ocular data.

The experimental session lasted approximately 2 hours and was comprised of two stages:

#### (1) Baseline ocular measures

Prior to the main experimental session, we performed a series of brief baseline measures of ocular reactivity. These included measuring responses to a slow, gradual change in screen brightness, to a sudden flashing white screen, to a sudden flashing black screen, and to a sudden presentation of a brief auditory stimulus (harmonic tone). These measurements are used to confirm normal ocular reactivity.

#### (2) Main experiment

In the main experiment, ocular data were recorded while participants listened to the artificial scene stimuli and performed the decoy gap detection task. A short practice was provided beforehand to ensure participants understood the task. Participants were naïve to the experimental conditions (scene changes and scene predictability) and were instructed to monitor for and quickly respond to the silent gaps. On trials on which a response was made (a correctly detected gap or a false alarm), feedback was provided. A summary of performance was also presented at the conclusion of each block. ∼3-minute breaks were provided between blocks. 8 blocks were completed.

All experimental tasks were implemented in MATLAB and presented via Psychophysics Toolbox Version 3 (PTB-3).

#### Procedure

Participants sat with their head fixed on a chinrest in front of a monitor (24-inch BENQ XL2420T with a resolution of 1920×1080 pixels and a refresh rate of 60 Hz) in a dimly lit and acoustically shielded room (IAC triple walled sound-attenuating booth). They were instructed to continuously fixate on a black cross presented at the centre of the screen against a grey background. An infrared eye-tracking camera (Eyelink 1000 Desktop Mount, SR Research Ltd.) placed below the monitor at a horizontal distance of 62cm from the participant was used to record eye data. Auditory stimuli were delivered diotically through a Roland Tri-capture 24-bit 96 kHz soundcard connected to Sennheiser HD558 headphones. The loudness of the auditory stimuli was adjusted to a comfortable listening level for each participant. The standard five-point calibration procedure for the Eyelink system was conducted prior to each experimental block, and participants were instructed to avoid any head movement after calibration. During the experiment, the eye-tracker continuously tracked gaze position and recorded pupil diameter, focusing binocularly at a sampling rate of 1000 Hz. Participants were instructed to blink naturally during the experiment and encouraged to rest their eyes briefly during inter-trial intervals. Prior to each trial, the eye tracker automatically checked that the participants’ eyes were open and fixated appropriately; trials would not start unless this was confirmed.

### Analysis of behavioural data

Gap detection task: Key presses that occurred <2 seconds following a target gap, were designated as hits. We also tracked the number of false alarms (responses when no gap was present) and reaction times (recorded from each hit). Related-Samples Wilcoxon Signed Rank test was conducted to test the main effect of regularity (REG and RND) on gap detection. Overall, participants made few false alarms; therefore, only hit rate and reaction time were analysed.

### Pupillometry Preprocessing and Analysis

Trials containing a gap or where a participant made a false alarm were excluded from the analysis. Where possible, data from the left eye were analysed. Intervals where the participant gazed away from fixation (outside of a radius of 100 pixels around the centre of the fixation cross) or where full or partial eye closure was detected (e.g. during blinks) were automatically treated as missing data and recovered using shape-preserving piecewise cubic interpolation. Data were then smoothed with a 150-ms Hanning window.

For NC trials, the pupil data were epoched from 1 s before stimulus onset to 1-sec post offset (−1: 10 sec). For change trials (CA and CD), the pupil data were epoched from 0.5 s before change time to 2.5 s after change time (−0.5: 2.5 sec). Epochs with more than 50% missing data or those determined to be particularly noisy where 10% or more of the data were identified as outlying (>3SD from the condition mean) were discarded from the analysis. On average, <6 trials were discarded per subject for all conditions. To allow for comparison across trials and subjects, data for each subject in each condition and each block were normalized. To do this, the mean and standard deviation across all baseline samples (1 s for NC and 0.5 s for CA and CD) in that block were calculated for each condition and used to z-score normalize the relevant epoched data. For each participant, pupil diameter was time-domain averaged across all epochs to produce a single time series per condition.

### Pupil dilation rate analysis

The pupil events were extracted from the continuous, smoothed data (150-ms Hanning window). Based on Joshi et al. (2016) and Zhao et al. (2019a; 2024), the events were defined as local minima (dilations; PD) that are followed by continuous dilation of the pupil for at least 100 ms. In each condition, for each participant, the event time series were summed and normalized by the number of trials and the sampling rate. Then, a causal smoothing kernel ω (τ) = α ^2^ × τ × *e*^−ατ^ was applied with a decay parameter of 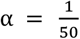 (Dayan and Abbott, 2005; Rolfs et al., 2008; Widmann et al., 2014) paralleling a similar technique for computing neural firing rates from neuronal spike trains (Dayan & Abbott, 2005; see also Joshi et al., 2016; Rolfs et al., 2008). The mean across trials was computed and baseline corrected. To account for the time delay caused by the smoothing kernel, the time axis was shifted by the latency of the peak of the kernel window.

### Microsaccade Preprocessing and Analysis

Intervals, where full or partial eye closure was detected (e.g. during blinks) were automatically treated as missing data and not interpolated. Microsaccade (MS) detection was based on an approach proposed by Engbert and Kliegl (2003). MS were extracted from the continuous eye-movement data based on the following criteria: (1) A velocity threshold of λ = 6 times the median-based standard deviation within each condition (2) Above-threshold velocity lasting between 5ms and 100ms (3) The events are detected in both eyes with onset disparity <10ms (4) The interval between successive microsaccades is longer than 50ms. Extracted microsaccade events were represented as unit pulses (Dirac delta). For NC trials, the data were epoched from 0.5 s before stimulus onset to 1-sec post offset (−0.5: 10 sec), and for change trials (CA and CD), the data were epoched from -0.15 s before change time to 2.5 s after change time (−0.15: 2.5 sec). Epochs with more than 50% missing data were discarded from the analysis. The microsaccade rate was then computed in the same way as described for the pupil dilation incidence rate, above.

### Statistical analysis

To identify time intervals showing significant PD/MS differences between conditions, we employed a non-parametric, bootstrap-based statistical analysis (Efron and Tibshirani, 1994). For each participant, we computed the difference time series between conditions and subjected these to bootstrap resampling (1,000 iterations with replacement). At each time point, differences were considered significant if more than 95% of the bootstrap iterations fell consistently above or below zero. This analysis was conducted across the full epoch.

In the analysis of sustained PD/MS effects and phasic pupil responses, we controlled for false discoveries using a permutation approach. Specifically, condition labels were randomly shuffled, and the same bootstrap procedure was applied. This process was repeated 500 times to generate a distribution of the longest significant intervals expected by chance. From this, we identified a 5% threshold for cluster length, which served as the significance criterion for the main bootstrap analysis—only clusters exceeding this threshold were considered significant. Given the transient nature of MSI effects, we did not apply false discovery rate (FDR) correction. Instead, any significant effects observed outside the predefined region of interest (100–250 ms, corresponding to typical MSI latency) were treated as exploratory.

## Results

### Gap detection performance

As expected, the decoy gap detection task was easy (Figure 2). Hit rates were high and close to ceiling in most participants. False alarm rates were very low across participants (<5 false alarms per subject). Pairwise Wilcoxon signed-rank tests were conducted to compare Hit rate and reaction time data across conditions. There was no difference between REG and RND conditions in terms of hit rate (Z = -1.458 p=.145) or reaction time (Z = -1.228 p=.219).

**Figure 2:**
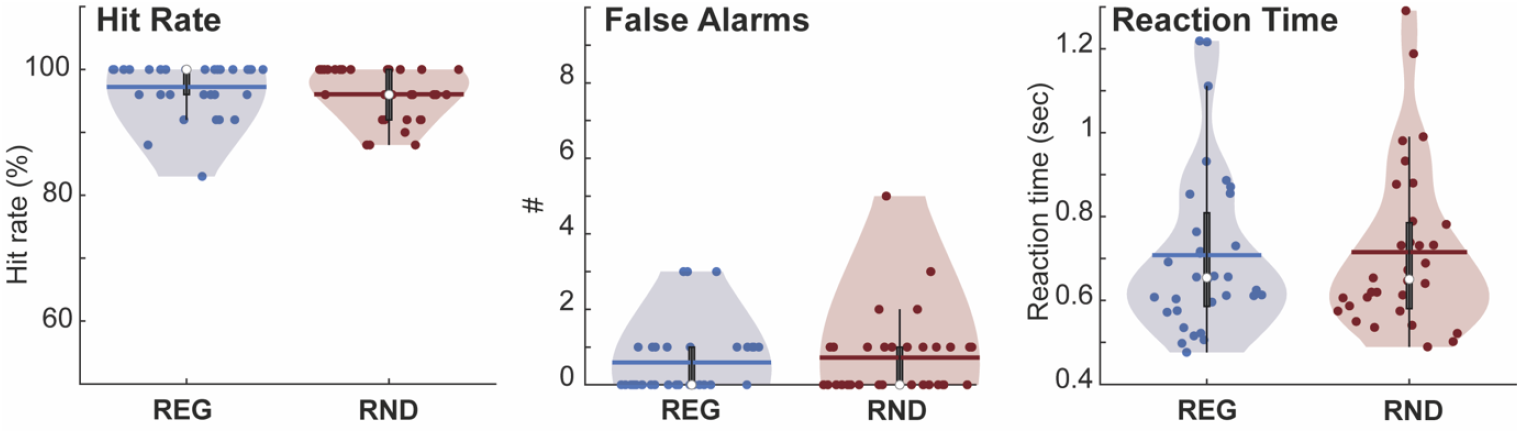
Decoy (Gap detection) task performance. Dots represent individual data. Error bars show SEM. No difference was observed in terms of RTs and hit rate between REG and RND conditions.

### PD and MS data reveal re-orienting responses to unattended scene changes

Figure 3 shows PD responses to scene changes against the no-change control (NC). Responses to both CA and CD (in both REG and RND scenes) show a pronounced increase in pupil diameter, consistent with a phasic response. The dynamics of these responses mirror dynamics observed in MEG data (Sohoglu and Chait, 2016b) a sharp early double-peaked response evoked by CA and a later, single-peaked response evoked by CD.

**Figure 3:**
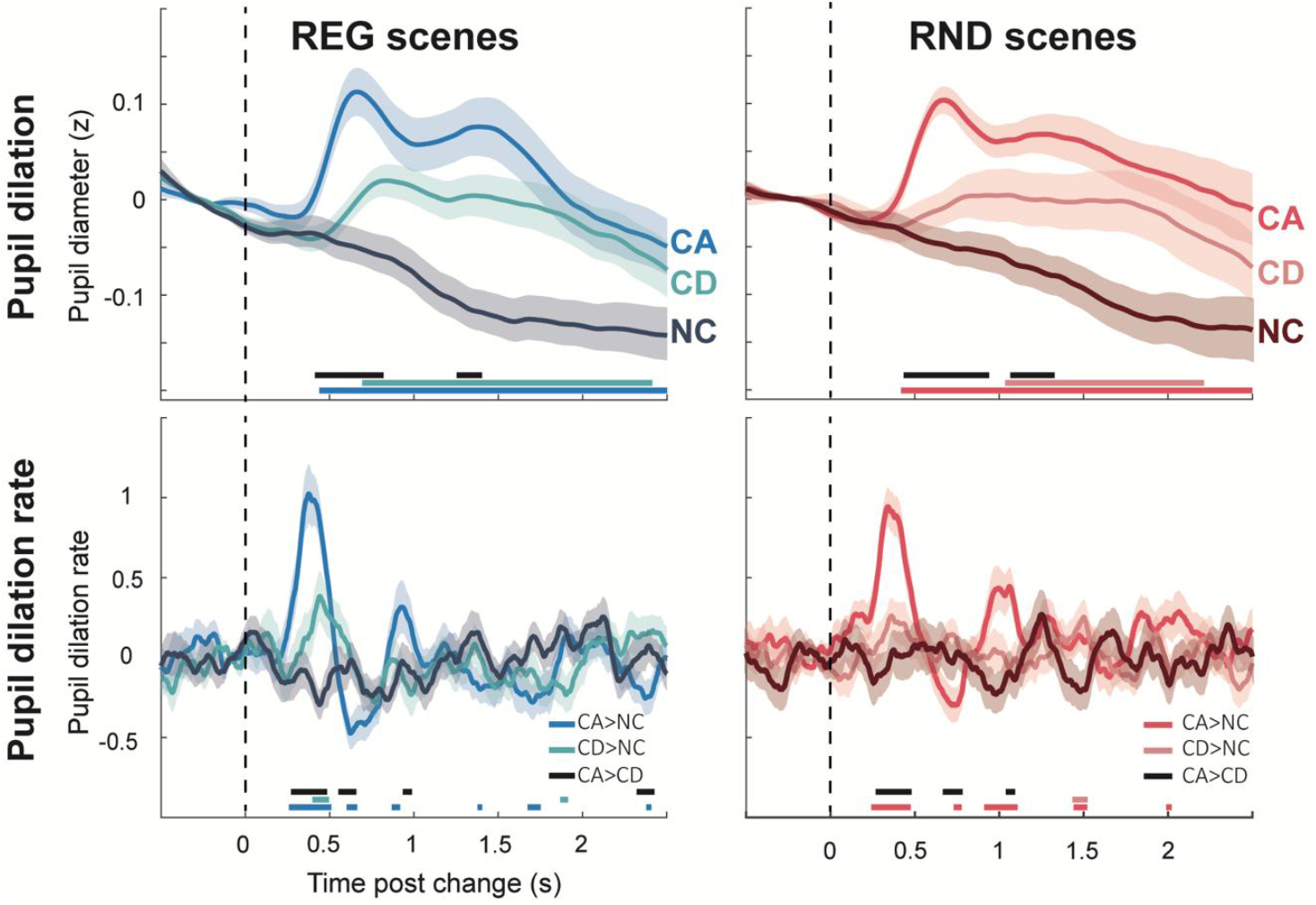
Change-evoked pupil dilation (top) and pupil dilation rate (bottom) across conditions (CA, CD, NC). Baseline corrected (–0.5 s to 0 s pre-onset for PD and -0.15 s to 0 s pre-onset for PDR). Shading indicates SEM. The horizontal bar indicates significant differences between conditions (p < 0.05; see legend below the bottom traces).

We also specifically focused on pupil-dilation events (see methods), as a potentially more sensitive measure of phasic pupil activity, which is associated with corresponding phasic activity in the Locus Coeruleus (Joshi et al. 2016; see also Jagiello et al, 2019 ; Zhao et al, 2024). These results revealed a consistent pattern, with a larger and earlier increase in pupil dilation rate evoked by CA events.

MS data are shown in Figure 4. A clear MSI response is visible for CA changes, though substantially larger in REG relative to RND scenes. For CD changes, a first difference between conditions emerges much later, at around 0.5 seconds post-onset. This is only seen in REG scenes, no differences are observed in RND scenes.

**Figure 4:**
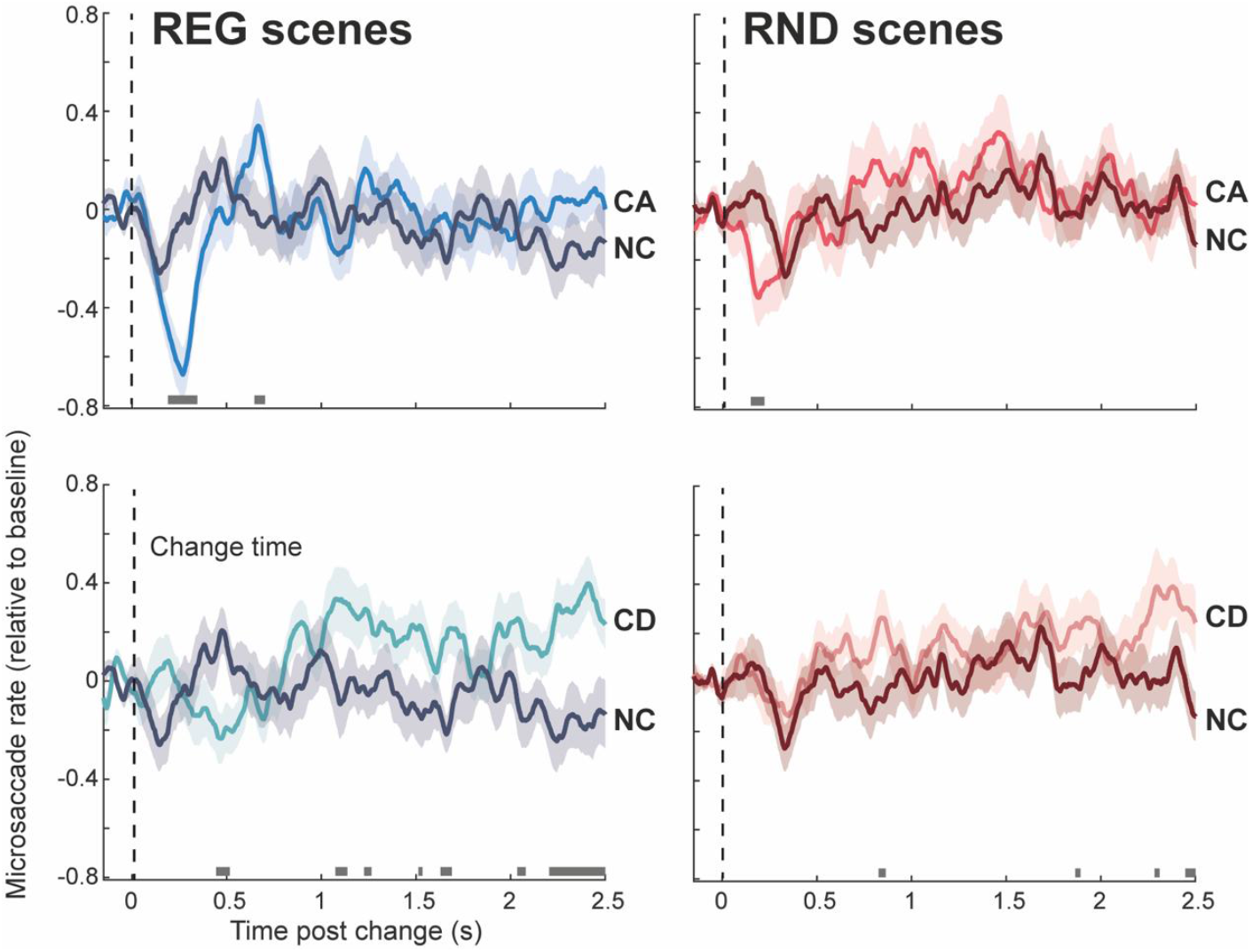
Change-evoked MS responses. Baseline corrected (−0.15s to 0-s pre-onset). The shaded area shows SEM. Horizontal bars indicate significant differences (p<0.05) between condition pairs.

Of interest is the apparent emerging sustained difference between CD and NC changes whereby the MS rate in CD trials exhibits a sustained rise over the NC rate. This incidental finding is difficult to interpret given the relatively weak significance but might indicate increased scene exploration in CD scenes, as discussed further below.

### Tonic PD reveals sensitivity to scene regularity

Figure 5 shows pupil responses to REG and RND scenes (no change). Both conditions revealed a pupil diameter increase shortly after scene onset, reaching an initial peak at 0.8. s post-onset, followed by a broader peak around 3 s after onset. Thereafter, the response entered a sustained phase, which lasted until the scene offset and was associated with a slow, continuous decrease in pupil diameter. Responses to REG and RND scenes overlapped initially but diverged after 3-sec post-onset, with REG scenes characterized by a faster decrease in pupil size than RND scenes. This pattern is consistent with what was previously shown by Milne et al. (2021) for regularly repairing vs random pip patterns. The results thus demonstrate that listeners’ arousal level, as reflected by pupil size change, is modulated by the regularity of the complex auditory scene.

**Figure 5:**
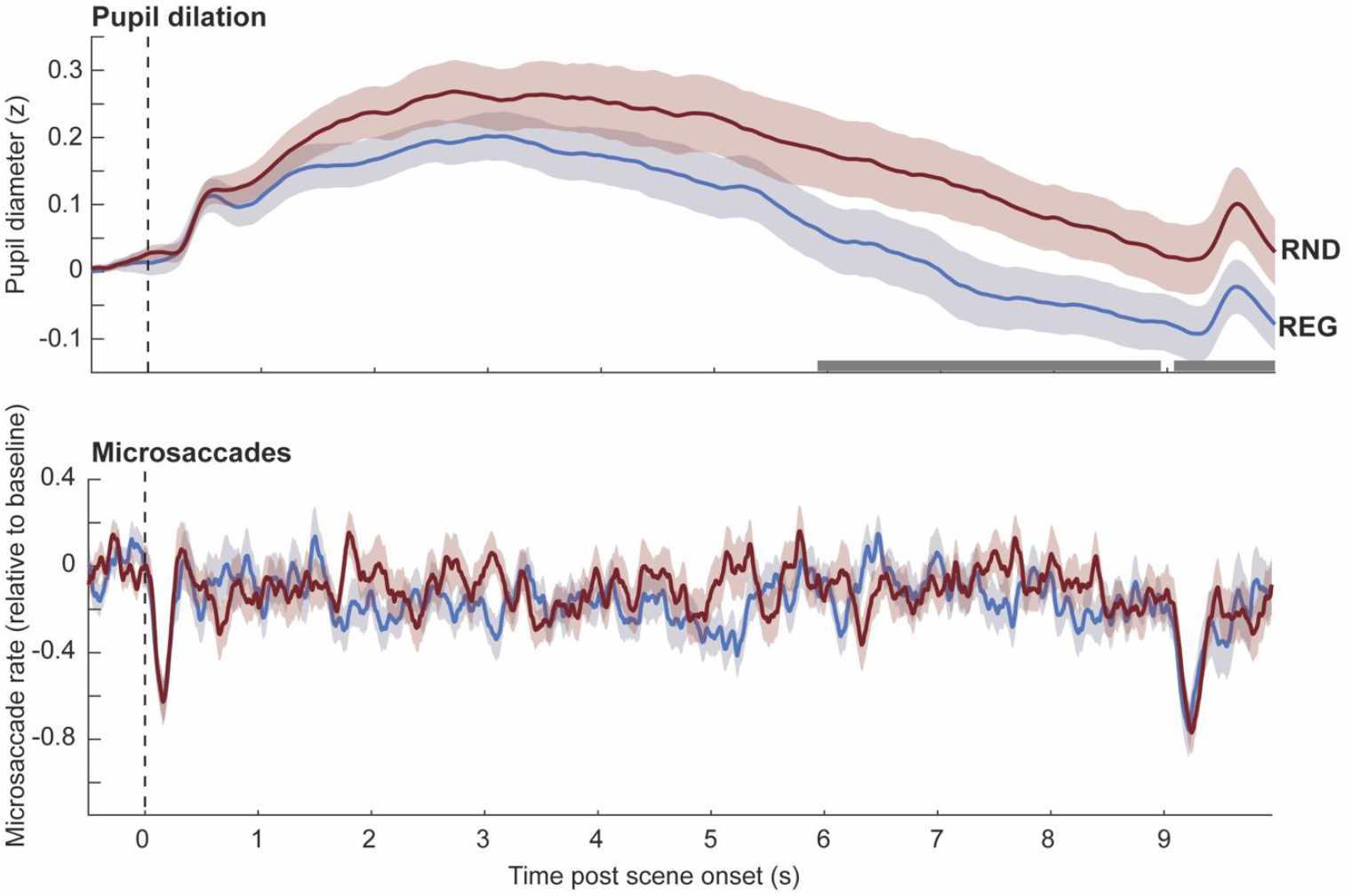
Sustained PD and MS responses to REG vs RND scenes. The shaded area shows SEM. The horizontal bars show time intervals during which significant differences were observed between conditions (p<0.05).

A similar analysis on MS data (Figure 5, bottom) did not reveal any differences between conditions. Clear MSI are seen following sequence onset and offset but the two conditions do not exhibit systematic differences.

### MS, but not pupil responses to unattended appearing and disappearing events are modulated by scene regularity

Figure 6 plots PD, PDR and MS responses to CA (left) and CD (right) changes in REG relative to RND scenes. Since REG and RND scenes are characterized by different sustained (tonic) pupil dilation baselines (see Fig 5 above), change-evoked responses were quantified by subtracting PD responses to CA/CD from the PD response to the control (no change; NC) condition. PD responses to both change types did not differ between REG and RND scenes. Similarly, no effect of regularity was observed in the PDR responses.

**Figure 6:**
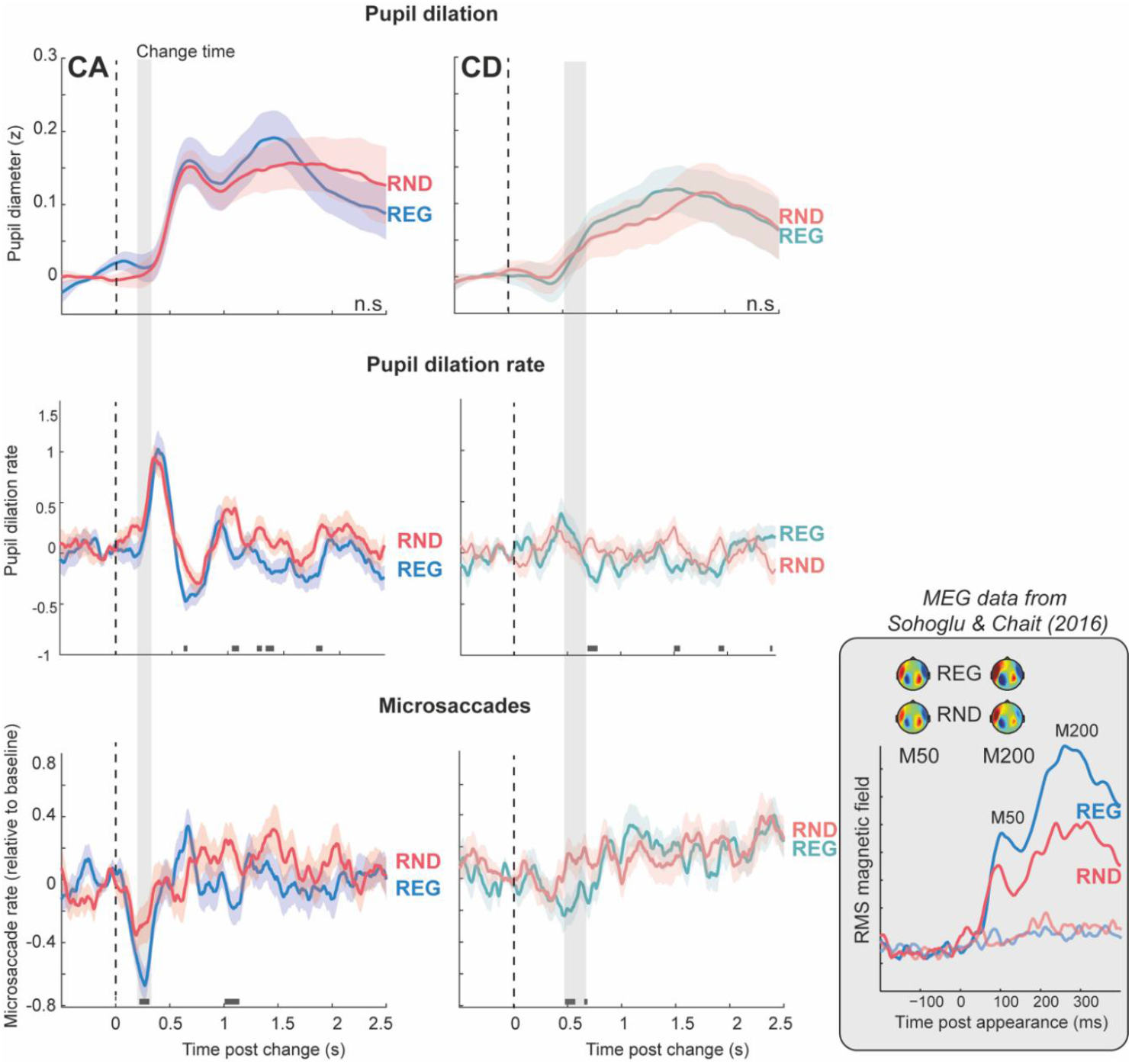
Effect of scene regularity on change-evoked MS, PD and PDR responses. Shading around the traces indicates SEM. Horizontal bars indicate intervals where a significant difference (p<0.05) is present between condition pairs. Grey shading marks the MSI effects in CA and CD scenes, highlighting the absence of corresponding differences in PD or PDR responses. The inset shows MEG data adapted from Sohoglu & Chait (2016a), illustrating enhanced responses to CA in REG compared to RND scenes from approximately 70 ms post-onset. The latency of the MSI effect for CA aligns broadly with the timing of these MEG effects.

In contrast, MS responses to CA/CD demonstrate a clear effect of regularity. CA changes in REG scenes are associated with a larger MSI response (more microsaccadic inhibition) than RND scenes. Significant differences between conditions emerge between 190-340 ms post-change onset. The MSI response appears to be triggered at the same time in both conditions but reaches a substantially lower trough in REG, indicative of a stronger attentional capture. The timing is similar to that observed in Zhao et al. (2024). An additional significant interval is observed between 1-1.3 s potentially reflecting that the appearing source in REG scenes—typically more perceptually salient—captured greater attention than in RND scenes.

A difference between REG and RND conditions was also observed for CD scenes, though it emerged substantially later than for CA—approximately between 0.47 and 0.68 seconds. This effect was accompanied by a more pronounced MSI response to CD in REG scenes. However, as noted above, the MSI response to CD is generally weaker and occurs later compared to that for CA.

## Discussion

Unattended changes in complex acoustic scenes elicit increased arousal and attentional capture, reflected in phasic pupil dilation and MSI. Sustained pupil dilation was reduced in regular (REG) compared to random (RND) scenes, suggesting that greater predictability is linked to lower arousal. Predictability also modulated MSI, with stronger attentional capture in REG scenes, though phasic pupil responses were unaffected.

Taken together, these findings highlight a dynamic interplay between arousal, as reflected in PD, and attentional capture, as indexed by MS, during auditory scene analysis.

### Pupil response dynamics to CA and CD mirror previously observed MEG responses

Pupil dilation (PD) is a well-established proxy for activity in the locus coeruleus– norepinephrine (LC-NE) system, which regulates arousal and attention (Aston-Jones & Cohen, 2005; Sara & Bouret, 2012; Joshi et al., 2016; Joshi & Gold, 2020). Phasic PD responses to unexpected events are thought to reflect transient LC-NE activity (e.g., Bala & Takahashi, 2000; Wang et al., 2014). Here, we show that unattended auditory changes—source appearances (CA) and disappearances (CD)—evoke such responses. The response to CD is slower and smaller than to CA, mirroring previous MEG findings (Sohoglu & Chait, 2016b). This timing difference may arise from the increased computational demands involved in detecting disappearances. While detecting an appearance can rely on the onset of new energy in a previously inactive frequency band, detecting a disappearance requires ongoing monitoring and comparison to prior acoustic states (Yamashiro et al., 2009; Cervantes Constantino et al., 2012; Andreou et al., 2015). The current pupil data suggest that these perceptual asymmetries are also reflected in differences in phasic arousal.

Interestingly, the temporal profile of the pupil response closely resembles that of the MEG signal. The MEG response to CA exhibits two peaks—at approximately 40 ms and 96 ms post-onset—believed to reflect, respectively, the neural response to the transient acoustic event and subsequent processes such as recognition or attentional capture (Sohoglu & Chait, 2016b). The observation of similarly biphasic dynamics in the pupil response suggests that these distinct neural processes may have temporally dissociable effects on arousal.

### MSI dynamics reveal attentional capture by scene changes

Microsaccades (MS) have gained attention in auditory research, with evidence showing that MS rates are modulated by auditory attention (Widmann et al., 2014; Abeles et al., 2020; Contadini-Wright et al., 2023). Even early microsaccadic inhibition (MSI)—a rapid reduction in MS rate following abrupt sensory events (Rolfs et al., 2008; Rolfs, 2009; Hafed et al., 2021) —is influenced by higher-order auditory factors (Kadosh & Bonneh, 2022; Zhao et al., 2024). Zhao et al. (2024) found that MSI was stronger and longer for attended versus unattended sounds, suggesting that high-level auditory processing interacts with oculomotor control.

Here, we show that unattended auditory scene changes modulate MS activity, indicating bottom-up attentional capture. Unlike the strong MSI evoked by source appearance (CA), disappearance (CD) elicits a weaker, delayed MS response—consistent with reduced behavioural sensitivity to CD (Cervantes-Constantino et al., 2012; Aman et al., 2021). A gradual increase in MS rate for CD relative to NC also emerges later in the trial, particularly in REG scenes, potentially reflecting an effort to resolve complex scene changes. Behavioural evidence shows that even when listeners detect the disappearance of a source, they often fail to identify which source has disappeared (Cervantes-Constantino et al., 2012). These MS dynamics may therefore reflect an automatic, information-seeking response (e.g., Schneider et al., 2020), with attentional resources shifting toward visual exploration, resulting in increased MS activity.

### Scene regularity is associated with PD-indexed reduced arousal but not with MS-indexed attentional capture

Sustained pupil diameter was reduced in REG compared to RND scenes, consistent with previous findings by Milne et al. (2021) using tone sequences. This aligns with the hypothesis that predictable patterns ease processing demands, thereby lowering cognitive load and reducing arousal—as reflected in smaller pupil size. In REG scenes, listeners can likely anticipate upcoming events within each stream, facilitating more efficient processing. In contrast, the unpredictability of RND scenes places greater demands on cognitive resources. Notably, the pupil diameter difference between REG and RND scenes emerged relatively late—around 6 seconds after sequence onset—mirroring the timing reported by Milne et al. (2021). This delayed effect contrasts with earlier differentiation seen in M/EEG data (e.g., ∼400 ms post-onset; Sohoglu & Chait, 2016a), suggesting that pupil responses reflect the downstream impact of predictability on arousal rather than the initial detection of regularity.

Conversely, sustained microsaccade (MS) rates did not differ between REG and RND scenes. Given the established link between sustained MS activity and heightened cognitive engagement (Dalmaso et al., 2017; Lange et al., 2017; Xue et al., 2017; Abeles et al., 2020; Contadini-Wright et al., 2023), this null result suggests that scene regularity does not modulate attentional capture. This finding is particularly relevant to ongoing debates surrounding the role of predictability in guiding attention (Feldman and Friston, 2010; Zhao et al., 2013; Southwell et al., 2017; Press et al., 2020). If REG scenes are more attentionally demanding in a bottom-up manner, we would expect to see corresponding differences in MS activity, as observed in studies of stimulus-driven attention. The absence of such differences here implies that attentional engagement was comparable across both conditions.

Taken together, these findings suggest that while scene regularity reduces arousal and potentially liberates processing resources, it does not produce consistent changes in attentional allocation.

### MSI, but not PD, is affected by bottom-up auditory attentional capture

Consistent with the idea that sensitivity to statistical structure supports efficient interaction with the environment (Winkler et al., 2009; Bendixen et al., 2010; Bendixen, 2014), prior studies have shown that change detection is more effective in structured (REG) than in random (RND) environments —reflected in faster reaction times and higher d’ values (Aman et al., 2021; de Kerangal et al., 2021).

Consistently, in Sohoglu and Chait (2016a), using stimuli essentially identical to the ones used here, change-evoked neural responses were significantly stronger in REG scenes, (see Figure 6). These findings support the notion that the auditory system automatically constructs precise models of the acoustic environment based on statistical regularities. Violations of these models—i.e., unexpected events—generate prediction errors, which elicit stronger neural responses and enhance perceptual salience (see also Garrido et al., 2013; Southwell & Chait, 2018; SanMiguel et al., 2021).

We observed phasic pupil responses to changes in both REG and RND scenes, but unlike the MEG data, did not find evidence of a “regularity advantage” in the pupil response. While null effects should be interpreted with caution, the pattern of results suggests that any effect of regularity on PD is minimal. Notably, prior work investigating attention-related effects on PD (Zhao et al., 2024) found robust effects with a comparable sample size, indicating that regularity-related effects—if present—should have been detectable in the current dataset. This lack of modulation in the PD data implies that the “predictability advantage” observed in behaviour and neuroimaging may not be mediated by changes in arousal.

In contrast, a clear effect of regularity was observed in the MS data. CA and CD events in REG scenes were associated with stronger MSI, indicating heightened attentional capture by changes in predictable contexts. Notably, the timing of this modulation—emerging subsequent to the initial sharp drop in MS incidence—closely matches the pattern reported by Zhao et al. (2024) for top-down attention. This reinforces the hypothesis that the earliest phase of MSI is not modulated by attention, and that attentional effects emerge at or after the MSI trough—an observation that can help constrain models of the underlying neural circuitry.

Interestingly, the timing of the MSI effect also aligns broadly with the MEG findings, where scene predictability influenced CA-evoked responses from ∼70 ms post-change under passive listening conditions. This suggests that the MEG and MS signals may reflect related components of a shared underlying process. In the MEG data, regularity effects were localized to a network including auditory areas in the superior temporal lobe and the left postcentral gyrus. It would be important for future work to elucidate the relationship between this auditory network and the MS-linked oculomotor systems.

### Divergence between MS and PD results suggests interplay between arousal and attention during auditory change detection

The divergence between MS and PD effects underscores that these measures likely capture distinct facets of cognitive processing. For change detection, it is plausible that the influence of regularity—presumably tied to rapid prediction error processing—primarily manifests as attentional capture. This form of engagement may be sufficient to support behavioral adaptation without necessitating elevated arousal levels.

Conversely, at the whole-scene level, a dissociation was observed in the opposite direction: PD, but not MS, reflected sensitivity to scene regularity (as discussed above), suggesting that a regular context reduces arousal without inducing changes in attention. While further research is necessary to unpack the nature of this dissociation, the current findings contribute to growing evidence that PD and MS serve as complementary readouts of separate stages in scene analysis. Specifically, they appear to index dissociable effects of attention and arousal within the broader framework of automatic auditory scene analysis.

## Acknowledgements

MH is supported by a PhD studentship from the Turkish Ministry of education. This work was supported by a BBSRC project grant to MC. The funders had no role in study design, data collection, and analysis, decision to publish, or preparation of the manuscript.

